# Characterization of SARS-CoV-2 public CD4+ αβ T cell clonotypes through reverse epitope discovery

**DOI:** 10.1101/2021.11.19.469229

**Authors:** Elisa Rosati, Mikhail V. Pogorelyy, Anastasia A. Minervina, Alexander Scheffold, Andre Franke, Petra Bacher, Paul G. Thomas

## Abstract

The amount of scientific data and level of public sharing produced as a consequence of the COVID-19 pandemic, as well as the speed at which these data were produced, far exceeds any previous effort against a specific disease condition. This unprecedented situation allows for development and application of new research approaches. One of the major technical hurdles in immunology is the characterization of HLA-antigen-T cell receptor (TCR) specificities. Most approaches aim to identify reactive T cells starting from known antigens using functional assays. However, the need for a reverse approach identifying the antigen specificity of orphan TCRs is increasing.

Utilizing large public single-cell gene expression and TCR datasets, we identified highly public CD4^+^ T cell responses to SARS-CoV-2, covering >75% of the analysed population. We performed an integrative meta-analysis to deeply characterize these clonotypes by TCR sequence, gene expression, HLA-restriction, and antigen-specificity, identifying strong and public CD4^+^ immunodominant responses with confirmed specificity. CD4^+^ COVID-enriched clonotypes show T follicular helper functional features, while clonotypes depleted in SARS-CoV-2 individuals preferentially had a central memory phenotype. In total we identify more than 1200 highly public CD4+ T cell clonotypes reactive to SARS-CoV-2. TCR similarity analysis showed six prominent TCR clusters, for which we predicted both HLA-restriction and cognate SARS-CoV-2 immunodominant epitopes. To validate our predictions we used an independent cohort of TCR repertoires before and after vaccination with *ChAdOx1*, a replication-deficient simian adenovirus-vectored vaccine, encoding the SARS-CoV-2 spike protein. We find statistically significant enrichment of the predicted spike-reactive TCRs after vaccination with *ChAdOx1*, while the frequency of TCRs specific to other SARS-CoV-2 proteins remains stable. Thus, the CD4-associated TCR repertoire differentiates vaccination from natural infection.

In conclusion, our study presents a novel reverse epitope discovery approach that can be used to infer HLA- and antigen-specificity of orphan TCRs in any context, such as viral infections, antitumor immune responses, or autoimmune disease.

**Highlights:**

- Identification of highly public CD4+ T cell responses to SARS-CoV-2
- Systematic prediction of exact immunogenic HLA class II epitopes for CD4+ T cell response
- Methodological framework for reverse epitope discovery, which can be applied to other disease contexts and may provide essential insights for future studies and clinical applications

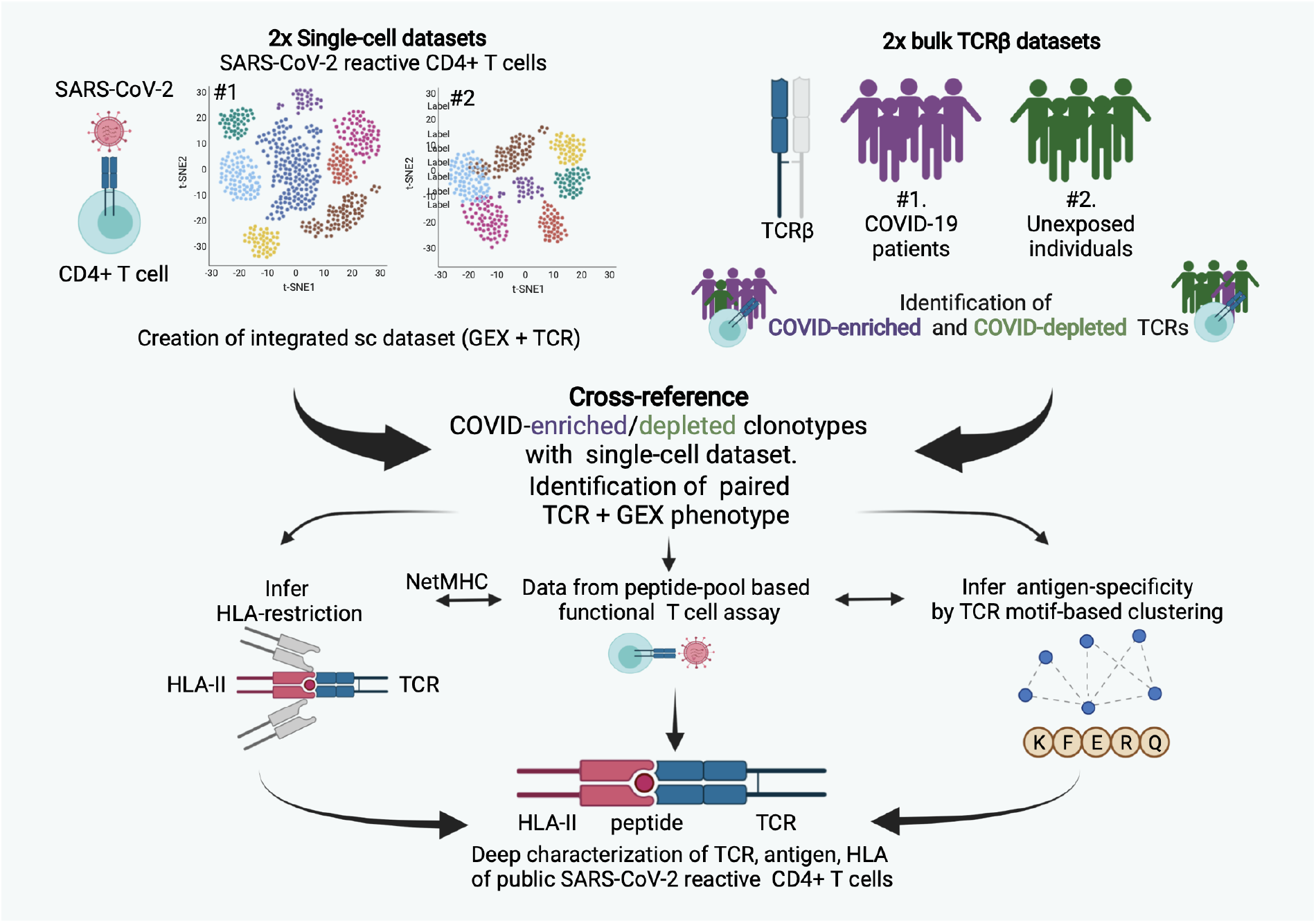

## Introduction

The worldwide scientific effort to overcome the COVID-19 pandemic led to the generation of an extraordinarily large amount of publicly available data describing the human immune response to SARS-CoV-2. A lot of these studies point to the importance of a T cell response in resolving COVID-19, as well as providing long-term protection against the variants of SARS-CoV-2 (Dan et al., 2021; Geers et al., 2021; Rydyznski Moderbacher et al., 2020; Sekine et al., 2020; Tarke et al., 2021a). One of the increasingly popular ways to study the complexity of the T cell response is T cell receptor repertoire sequencing (Mukhopadhyay, 2021). The T cell receptor (TCR) is a heterodimer of alpha and beta chains, both of which are formed in a semi-random DNA recombination process resulting in a unique repertoire in each individual (Dupic et al., 2021). However, even individual diverse T cell repertoires can show similar features after encountering the same antigens. In particular, response to immunodominant epitopes triggers large clonal expansions, and TCRs recognising such epitopes frequently have highly similar sequences (Dash et al., 2017; Glanville et al., 2017; Pogorelyy et al., 2019). Thus, analysis of TCR repertoires could shed light on the differences and commonalities of the T cell immune response among different individuals, on the identity of the most immunogenic antigens, and may provide targets for development of diagnostic tools as well as therapeutic treatments such as adoptive T cell transfers (NLM, NCT04762186 clinical trial). As a proof of principle, identification of T cell clonotypes reactive to SARS-CoV-2 antigens has eventually led to the development of a diagnostic test for SARS-CoV-2 authorized for emergency use by the FDA (Dalai et al., 2021).

In order to identify TCR repertoire signatures related to COVID-19, multiple groups utilized bulk TCR repertoire sequencing (Minervina et al., 2021a; Niu et al., 2020; Nolan et al., 2020; Schultheiß et al., 2020; Shomuradova et al., 2020; Snyder et al., 2020), that quantitatively measures frequencies of large numbers of unpaired TCRalpha or TCRbeta clonotypes, as well as single cell TCR sequencing techniques (Bacher et al., 2020; Bernardes et al., 2020; Kusnadi et al., 2021; Liao et al., 2020; Lu et al., 2021; Meckiff et al., 2020; Wen et al., 2020; Xu et al., 2020; Zhang et al., 2020), that produce many fewer, but paired, alpha/beta TCR sequences. One of the major challenges of such approaches is that, even at the peak of the immune response to SARS-CoV-2 infection, only a small fraction of total peripheral T cells recognize viral epitopes. Hence, many studies rely on T cell antigen-enrichment methodologies, such as MHC-multimer-staining or peptide stimulation with subsequent enrichment for activated cells (e.g. ARTE assay, AIM assay, MIRA assay) (Altman et al., 1996; Bacher et al., 2013; Klinger et al., 2015; Reiss et al., 2017). These approaches increase the number of SARS-CoV-2 specific TCRs detected in each sample and help to identify immunodominant epitopes (reviewed in Grifoni et al. 2021). Stimulating T cells with peptide libraries is the most frequent approach used for SARS-CoV-2 epitope-discovery (Braun et al., 2020; Grifoni et al., 2020; Lu et al., 2021; Mateus et al., 2020; Nelde et al., 2021; Peng, Yanchun et al., 2020; Tarke et al., 2021b). Here, we focus on public clonotypes (clones found in multiple individuals), which provide the necessary power for a robust statistical analysis and, in addition, hold the highest potential for further, population-wide applications. We propose a reverse epitope discovery technique, which, instead of starting from the epitopes to identify reactive T cells, utilizes TCR repertoires as the means to find immunodominant responses in an unbiased manner. We performed a comprehensive TCR meta-analysis of publicly available single cell and bulk TCR repertoire datasets and identified SARS-CoV-2 reactive TCRs with complete TCR alpha and beta chain information and inferred their HLA restriction. Moreover, through TCR clustering based on sequence similarity, we were able to identify several prominent alpha/beta TCR motifs and predict their antigen specificity.

## Results and discussion

In this report, we jointly analysed **(1)** two published single-cell datasets of SARS-CoV-2-reactive CD4^+^ T cells, identified based on CD154+ up-regulation after peptide pool stimulation and including gene expression and TCR information for a total of 59 individuals, of which 49 were COVID-19 patients and 10 healthy controls (Bacher et al., 2020; Meckiff et al., 2020) and **(2)** the largest published bulk TCRbeta datasets of 786 healthy individuals (Emerson et al., 2017) and 1414 COVID-19 patients (Snyder et al., 2020), including TCRs with known specificity for certain SARS-CoV-2 peptide pools (Nolan et al., 2020) and **(3)** a published bulk TCR dataset before and after SARS-CoV-2 vaccination with *ChAdOx1* (Swanson et al., 2021), which we used as validation for our findings.

In order to find public CD4^+^ T cell responses to SARS-CoV-2 infection we first merged two publicly available single-cell datasets of CD4^+^ SARS-CoV-2-reactive T cells (Bacher et al., 2020; Meckiff et al., 2020). Both datasets were obtained using the same antigen-reactive T cell enrichment assay (ARTE-assay), based on CD154+ up-regulation on antigen-reactive CD4+ T cells (Bacher et al., 2013). Of note, differently from Bacher et al. the antigenic pool used in Meckiff et al. only includes peptides from the spike protein (without N-terminal domain) and the membrane glycoprotein. The combined dataset contained 125,258 cells that passed quality control steps, which resulted in 13 functional clusters after unsupervised analysis (**Figure 1A**). Cluster phenotypes were defined using markers of cell populations used in the original publications (Bacher et al., 2020; Meckiff et al., 2020). In particular, we found clusters corresponding to helper follicular T (TfH) cells (clusters 1-2), helper type 1 T (Th1) cells (cluster 3), transitional Tfh/Tcm T cells (cluster 4), central memory T (Tcm) cells (clusters 5), Th17 phenotypes (clusters 6-7), effector memory T cells (cluster 8), type I IFN-signature T cells (clusters 9-10), cytotoxic T cells (clusters 11-12), and cycling T cells (cluster 13) (**Figure 1A-B, Supplementary Table 1**). Using this extended dataset, we also confirmed previous findings of a significant enrichment of TfH T cells (Clusters 1 and 2) in COVID-19 patients in comparison to healthy controls (**Figure 1C-D**). Interestingly, the abundance of the two main TfH T cell subsets was significantly different in hospitalized versus non-hospitalized COVID-19 patients (**Figure 1D**). In particular, cluster 2 was enriched in severe disease, and expressed higher cytotoxic markers such as *CCL3, CCL4, CCL5, XCL1, XCL2, GZMB* and *GNLY.* In contrast, cluster 1 expressed higher levels of *IL2, CD69* and genes of the *TNF* family, and was nominally enriched in mild-disease patients as indeed previously shown by Meckiff *et al.* (**Figure 1D, Supplementary Table 1**).

**Figure 1:**
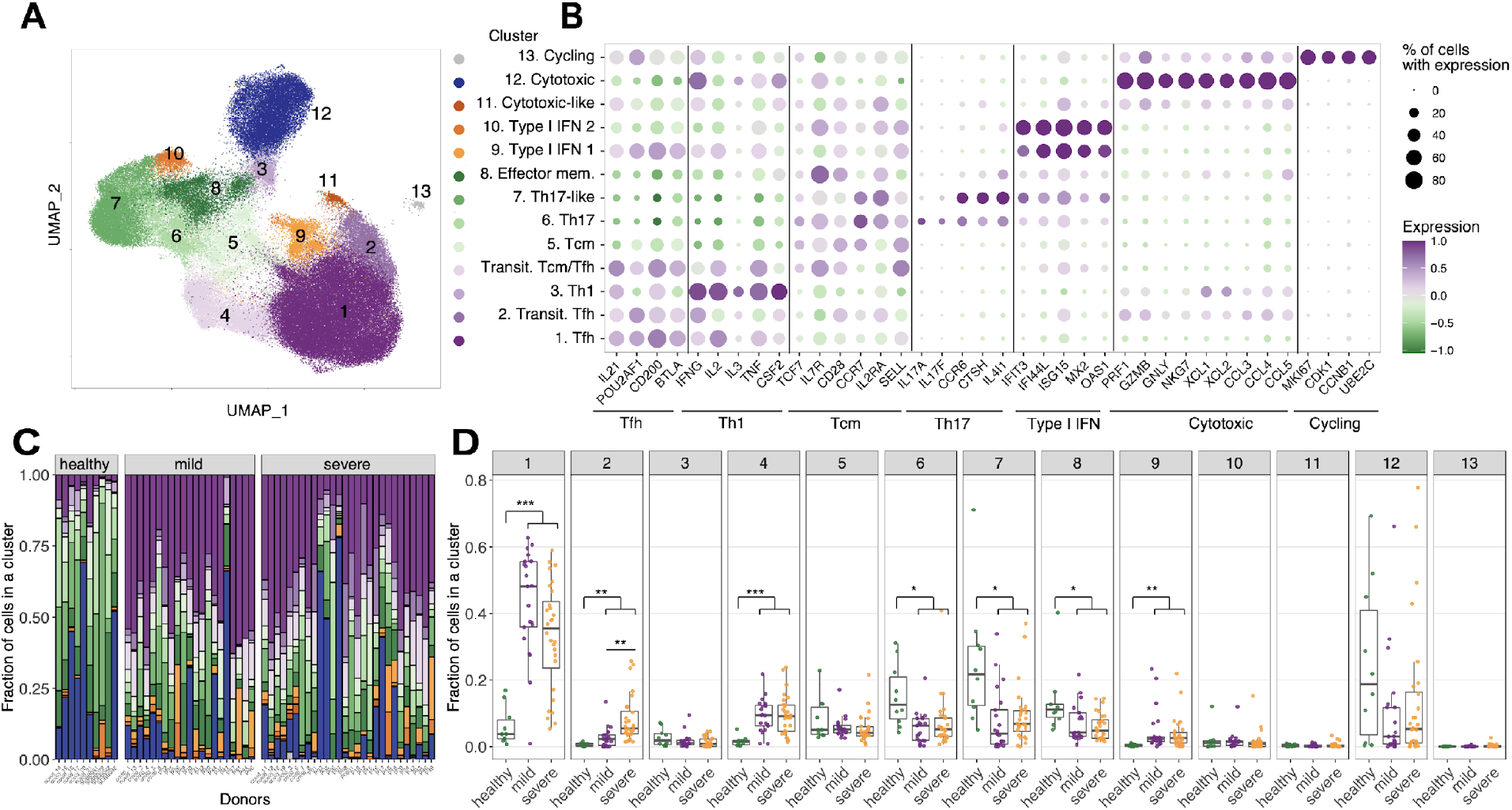
**A**. UMAP of single cells for merged (Bacher et al., 2020; Meckiff et al., 2020) datasets of SARS-CoV-2 antigen-enriched CD4 T cells based. Colors show clusters of cells with distinct gene expression profiles **B**. Differentially expressed genes in each GEX cluster. **C.** Distribution of cells between GEX clusters is plotted for each donor. Healthy donors less Tfh cells (populations 1 and 2) **D.** Boxplots showing cell proportion distribution among functional clusters for each patient (Mann-Whitney U test, Bonferroni multiple comparison correction).

In order to select TCRs corresponding to the most public CD4^+^ T cell responses, we next searched TCRbeta sequences from the combined single cell dataset in the TCR repertoires from a large cohort of COVID-19 patients (Snyder et al., 2020), as well as from pre-pandemic COVID19-naive controls (Emerson et al., 2017). We identified TCRs shared among individuals and strongly associated with SARS-CoV-2 infection, as defined by presence in significantly more patients than controls (COVID-enriched TCRs) (Fisher’s exact test), which are reported in **Supplementary Table 2**. We also identified TCRs which were significantly decreased in COVID-19 patients as compared to the healthy population (COVID-depleted TCRs), reported in **Supplementary Table 3** (**Figure 2A)**. Notably, when mapping COVID-enriched TCRs to the single-cell RNAseq data, these significantly accumulated in TfH T cell clusters (clusters 1,2,4) while COVID-depleted TCRs rather accumulated in effector memory subpopulations (clusters 6,7,8) (**Figure 2B**). The effect size (log2-fold enrichment) for COVID-depleted TCRs was much smaller than for COVID-enriched TCRs (**Figure 2A).** Moreover, COVID-depleted TCRs were present in a large fraction of donors from both the control and the COVID-19 cohorts (**Figure 2C)**, e.g., the majority of COVID-depleted TCRs (465 out of 594) were simultaneously found in >100 controls and >100 COVID-19 patients, while only 24 of 1248 COVID-enriched clonotypes had the same level of publicity. The number of unique T cell clones in a subset of the analyzed COVID-19 patients was low in comparison to healthy controls, potentially due to COVID-19 associated lymphopenia. This could lead to small, yet significant, underrepresentation of highly public clonotypes in the COVID-19 cohort. We therefore focused on the COVID-enriched TCR clonotypes for further analysis, because the occurrence pattern and phenotype of this group is consistent with expansion of T cell clones specific for SARS-CoV-2 antigens.

**Figure 2:**
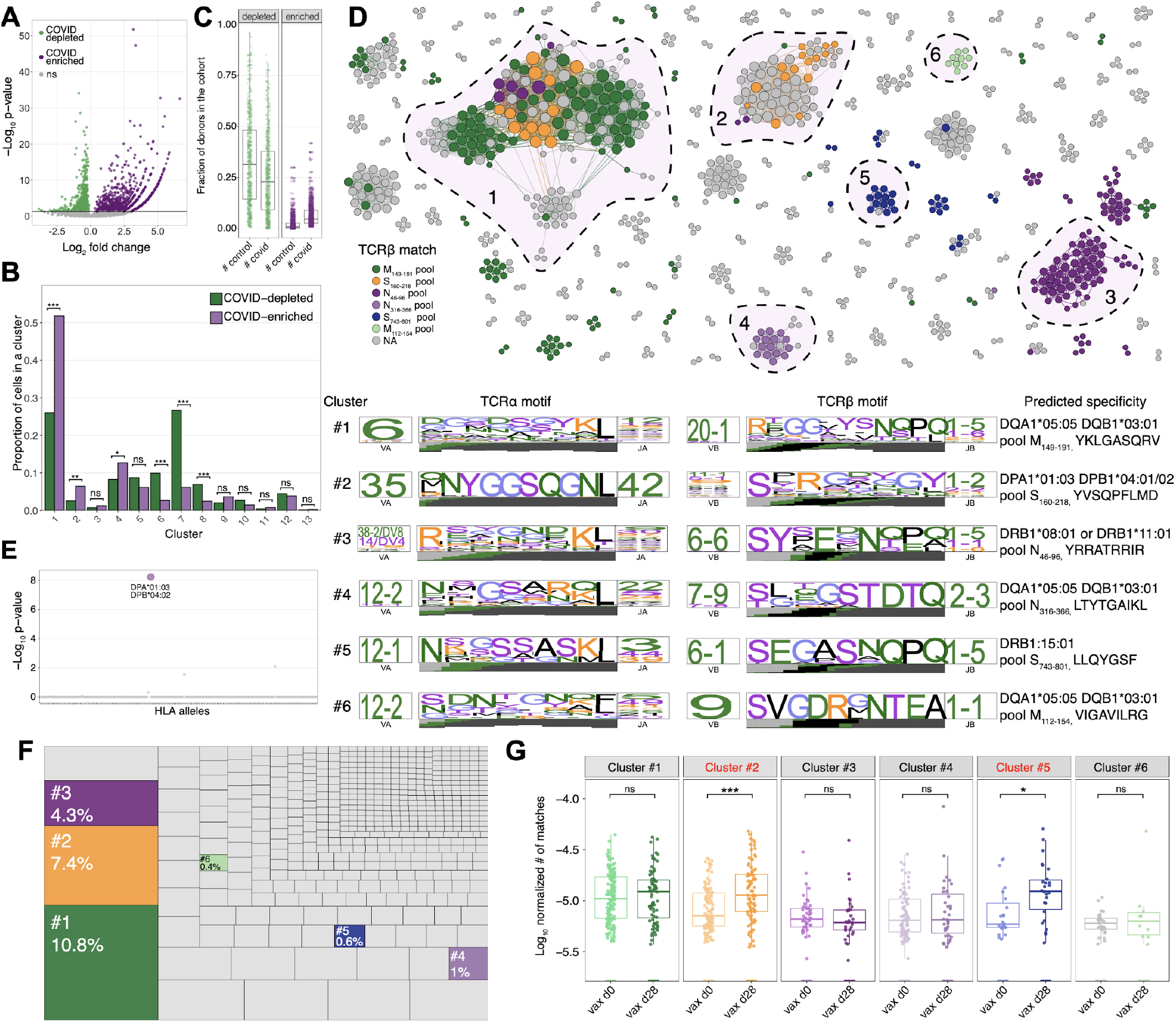
**A.** Volcano plot shows enrichment of TCRbeta chains identified in merged scTCRseq dataset in large (n=1414) collection of repertoires from COVID patients ((Snyder et al., 2020), purple) in comparison to the healthy donor cohort from Emerson et al (n=786) (x-axis) vs p-value (y-axis). **B**. Barplot showing the distribution of COVID-enriched (purple) and COVID-depleted (green) TCR clonotypes in GEX clusters. Fisher exact test was used for the comparison, with Bonferroni multiple comparison correction. **C.** The boxplots show a fraction of donors from healthy and COVID-19 cohorts sharing significantly COVID-depleted (green) and COVID-enriched (purple) clonotypes. **D.** Similarity network of COVID-associated public TCR clonotypes. Each vertex represents a TCR alpha/beta clonotype, edges connect vertices with <120 TCRdist units. Colors show predicted SARS-CoV-2 peptide pool from MIRA class II dataset. Bottom: TCRdist logos for the most prominent clonotype clusters with predicted peptide specificity and HLA-restriction. **E.** Manhattan plot for association of representative cluster #2 clonotype with various HLA-types. **F.** A treemap showing the fraction of repertoire occupied by clonotypes from prominent TCR similarity clusters from (c). **G.** Occurrence of TCRbeta from 6 large clusters from (c) before and after SARS-CoV-2 vaccination with ChAdOx1 vaccine. Significantly more TCRs from spike-specific clusters #2 and #5 are found after vaccination (one sided Wilcoxon rank sum test with Benjamini-Hochberg multiple testing correction).

Next, we aimed to identify TCRs with highly similar sequence motifs and thus likely to have the same antigen specificity (Dash et al., 2017; Glanville et al., 2017). We used TCRdist (Dash et al., 2017) to identify highly similar TCR sequences among TCRs enriched in COVID-19 and indeed found several prominent TCR clusters (**Figure 2D**). We also inferred the HLA-restriction for most of these TCRs, using the procedure suggested in (Minervina et al., 2021a), see **Figure 2E** for the representative Manhattan plot (and **Supplementary Table 2** for predicted HLA-restrictions).

TCRdist uses both TCRalpha and TCRbeta amino acid sequences to compute the distance between clonotypes. However, analysis of the resulting sequence motifs demonstrate unequal contribution of TCR alpha and beta chains in different clusters. For example, cluster 3 is largely defined by a conserved beta chain motif, allowing for diverse alpha chains, while in cluster 2 there is an almost invariant alpha chain paired with a set of very diverse TCRbeta chains. In a few other large clusters both TCR chains show strongly conserved amino acid motifs (**Figure 2D**). These differences could be potentially explained by the variable number of contacts of TCRalpha/TCRbeta chains with the antigenic peptide and MHC. Thus alpha-driven, beta-driven and alpha/beta driven motifs are interesting targets for solving TCR-pMHC ternary structures. To predict the potential antigen-reactivity of the TCR clusters we next cross-referenced the TCRbeta sequences of COVID-enriched clonotypes with the MIRA MHC-II dataset. This database contains specificity information of certain CD4^+^ TCRbeta sequences towards peptide pools of SARS-CoV-2 proteins. Specifically, in the MIRA MHC-II experiment, TCR-antigen specificity was tested against 56 peptide pools containing 1 to 6 overlapping peptide 19mers each, and spanning over the membrane (M), nucleocapsid (N) and spike (S) SARS-CoV-2 proteins. Our set of COVID-enriched TCRs mapped to 22 peptide pools, with most matches to M_149-191_, N_46-96_ and S_743-801_ pools. The number of TCRs matching into the MIRA database was significantly larger in the COVID-enriched TCRs (34%, 428 out of 1248) as compared to the COVID-depleted TCRs (13%, 80 out of 594) suggesting that the former group is indeed more enriched for common COVID-reactive clonotypes. Most TCRs belonging to the same TCRdist cluster were assigned to the same peptide pool from the MIRA MHC-II dataset.

Inference of cognate peptide pools and HLA-restriction allowed us to use NetMHC to predict specific antigenic peptides within the pool and also to validate our prediction. In fact, the predicted antigenic peptide and HLA restriction for TCRdist cluster 2 and cluster 5 exactly match the experimental results from (Mudd et al., 2021) and (Lu et al., 2021) respectively, thus supporting the validity of our methodology.

Finally, to validate the set of COVID-enriched CD4^+^ clonotypes using an independent dataset, we used a large collection of TCRbeta repertoires before and after SARS-CoV-2 vaccination with *ChAdOx1* (Swanson et al., 2021), a replication-deficient simian adenovirus-vectored vaccine, encoding the SARS-CoV-2 spike protein. For each TCR repertoire from the pre-vaccination (day 0) or post-vaccination time point (day 28) we calculate the fraction of unique TCRbeta clonotypes matching TCRbeta sequences from each of our largest antigen-specific TCR clusters (**Figure 2D**). We find significant enrichment of the predicted spike-reactive TCRs of clusters #2 and #5 after vaccination with *ChAdOx1*, while the frequency of TCR clusters reactive to the membrane or nucleocapsid proteins remain unchanged (**Figure 2G**). This result serves as an independent validation of our approach and, at the same time, shows how the TCR clusters we identified may be potentially used to identify SARS-CoV-2 epitope-specific TCRs both in the context of vaccination and natural infection.

## Conclusions

In conclusion, our study identified 1248 paired TCR clonotypes potentially specific to highly immunogenic epitopes from SARS-CoV-2. Many of these TCRs remain orphan of their epitopes. However, we identified and inferred antigen-specificity for 428 TCRs with the aid of the MIRA dataset (Nolan et al., 2020). We also inferred HLA-restriction for most of these TCRs. TCR-HLA pairings were also validated based on NetMHC binding predictions and functional experiments in other publications (Lu et al., 2021; Mudd et al., 2021) The resulting set of highly characterized public TCRs reactive to SARS-CoV-2 covers more than 76 percent of individuals from Snyder et al, i.e. at least 20 unique COVID-enriched TCR sequences per individual were found. The broad coverage of HLA haplotypes in our study also provides means to further investigate immunodominant responses in a genetically diverse population.

No immunodominant SARS-CoV-2 epitopes had yet been reported for most HLA class II alleles, especially if underrepresented in European populations. Furthermore, the high publicity of the characterized clonotypes makes them promising candidates for further studies on CD4+ T cell immunity against SARS-CoV-2 as well as for immunotherapeutic applications aimed at utilizing highly specific, immunodominant, T cell responses in the context of precision and personalized medicine (Müller et al., 2021; O'Reilly et al., 2016; Qian et al., 2018).

### Study limitations

The described method of reverse epitope discovery also has limitations that we want to highlight here. The method is strongly focused on public T cell responses which, on one side, make the findings significant and applicable to a larger fraction of the population. On the other side, immunodominant responses may be also driven by private clonotypes or clonotypes without an identifiable motif cluster, which would not be detected by our method (or that would not appear among the most interesting hits). Although this is a limitation of our approach, it may be a field to expand to in order to identify more T cell responses. Similarly, the publicity of HLA alleles is also a limiting factor of our method, as clonotypes recognizing peptides presented on rare HLA alleles would be hard to detect.

Another limitation lies in the availability of TCR antigen-specificity information from public databases and resources, such as VDJdb (Bagaev et al., 2020) and the MIRA dataset (Nolan et al., 2020). The expansion of these resources is of utmost importance for target identification of orphan TCRs. The strong general interest in the COVID-19 pandemic led to an unprecedented production of high amounts of TCR repertoire data, which is until now unmatched for other antigens or diseases.

Despite these limitations, we think that the reverse epitope discovery approach already proved itself valuable for identifying both cross-reactive (Minervina et al., 2021b) and immunodominant responses to SARS-CoV-2 (Mudd et al., 2021) and it holds potential for application in other disease contexts.

## Methods

### Utilized public data

Single-cell data of SARS-CoV-2 reactive CD154^+^ T cells were obtained from (Bacher et al., 2020) and (Meckiff et al., 2020). Bulk TCR data of healthy individuals and COVID-19 patients were the datasets used in (Emerson et al., 2017) and (Snyder et al., 2020), respectively.

### Single-cell datasets integration and filtering

The preprocessing of the scRNAseq data was performed with the 10x Genomics’ Cell Ranger software v3.1.0 using the human genome reference GRCh38 v3.0.0 for the mappings. The resulting raw feature-barcode matrix files were analyzed with the R package Seurat v.3.2.0 (Butler et al., 2018). Thereby, all genes with a detected expression in less than 0.1% of the non-empty cells were excluded. Moreover, TCR genes were not considered for further analyses to avoid functional clustering of cells based on TCR information. To minimize the number of doublets, empty cells, and cells with a low-quality transcriptome, only cells harboring between 400 and 3000 RNA features and less than 5% mitochondrial RNA were selected for further processing.

TCR information was integrated into the metadata of the Seurat object after filtering of cells containing more than 2 TCR alpha or 2 TCRbeta chains. After merging of Seurat metadata and TCR information, cells without TCR information were excluded from further analysis. Afterwards, data were log-normalized and scaled based on all genes. After performing a PCA dimensionality reduction (40 dimensions) with the RunPCA function, the expression values were corrected for batch effects caused by different sources of the data, sample preparation batches and sequencing run batches using the R package Harmony v1.0 (Korsunsky et al., 2019). In the final steps, the Uniform Manifold Approximation and Projection (UMAP) dimensional reduction was performed with the RunUMAP function using 40 dimensions, a shared nearest neighbor graph was created with the FindNeighbors method, and the clusters identification was performed with a resolution of 0.4 using theFindClusters function. 13 clusters were identified. Cluster marker genes were determined using FindMarkers with the MAST method (Finak et al., 2015).

### COVID-19 TCR association using bulk TCR public datasets

To identify public TCRbeta clonotypes we used two large datasets, one of COVID patients (n=1414, Snyder et al. 2020) and one of healthy subjects sampled pre pandemic (n=786, Emerson et al. 2017). For each TCRbeta from combined single cell TCRseq dataset we calculate number of unique donors from both bulk TCRbeta repertoire cohorts sharing it (a TCRbeta is considered shared if both CDR3 amino acid sequence and V segment family match, as suggested in (DeWitt et al., 2018)). Next, we use a two-sided Fisher exact test with Benjamini-Hochberg multiple-testing correction to identify sequences overrepresented (i.e. found in more donors) in either cohort (adjusted p-value<0.05 is used as significance threshold).

### Identification of motifs in TCR amino acid sequences using TCRdist

We used the TCRdist implementation in *conga* python package to calculate pairwise TCRdist between unique abTCR sequences and plot sequence logos for TCR motifs (Schattgen et al., 2021). We define TCR motifs as connected components on the TCR similarity network, where each node is a unique alpha/betaTCR clonotype, and edge connects them if distance is less than 120 TCRdist units. To filter TCR chimeras and other artifacts occuring during 10x Genomics sequencing resulting in rare spurious connections between motif clusters, we deleted top 1% of nodes and vertices by network betweenness centrality values. *igraph R* package was used to manipulate similarity networks (Csardi and Nepusz, 2006), *gephi* was used for network layout and visualisation (Jacomy et al., 2014).

### HLA specificity imputation from TCR data

For each donor from (Emerson et al., 2017; Snyder et al., 2020) we use HLA-types inferred in (Minervina et al., 2021a). Then for each TCRbeta significantly enriched in COVID cohort we do a one-sided Fisher exact test with Benjamini-Hochberg multiple testing correction to check if given TCRbeta co-occurs with each of HLA-alleles.

### Prediction of COVID-enriched TCR specificity

We mapped TCRbeta chain sequences from aggregated single cell dataset to peptide-pool specific TCRbeta clonotypes for MIRA class II dataset (release 002.1) allowing for one amino acid mismatch in CDR3 amino acid sequence. Next we selected six large clusters on the TCRdist similarity network with distinct MIRA peptide pool assignments, calculated consensus MIRA pool and HLA-restriction within a cluster and used NetMHCIIpan-4.0 (Reynisson et al., 2020) to predict the antigenic peptide within peptide pool.

### Statistical analysis

Statistical analysis was performed in R version 4.0.2. Wilcoxon rank-sum test (Mann-Whitney U test) was used to compare the proportion of cells in each Seurat functional cluster between healthy controls and COVID-19 patients as well as between severe and mild COVID-19 cases. Fisher exact test was used to compare the number of COVID-depleted and COVID-enriched clonotypes being part of each functional Seurat cluster. Multiple testing correction was performed using the Benjamini-Hochberg procedure. Ns not significant, * p<0.5, **p<0.01, ***p<0.001

## Data and code availability

All analysed data are publicly available. In detail, single-cell data can be found under SRA accession numbers SRP293741 (Bacher et al., 2020) and SRP267404 (Meckiff et al., 2020). Bulk TCR repertoire data from COVID infected subjects, and MIRA Class II dataset (release 002.1) (Snyder et al., 2020) are publicly available from ImmuneAccess database (https://clients.adaptivebiotech.com/pub/covid-2020), as well as healthy control data from (Emerson et al., 2017) (https://clients.adaptivebiotech.com/pub/emerson-2017-natgen) and *ChAdOx1* immunized cohort (https://doi.org/10.21417/PAS2021STM). Utilized scripts for the analysis of the merged single-cell dataset are available on the GitHub page https://github.com/pogorely/reverse_epitope_discovery

## Acknowledgements

This research was supported by the German Research Foundation (DFG) under Germany’s Excellence Strategy—EXC 2167-390884018 Precision Medicine in Chronic Inflammation to P.B., A.F., and A.S.; DFG 433038070 to P.B., A.F., and A.S.; DFG 4096610003 to E.R; by a COVID-19 research grant from the Land Schleswig-Holstein, DIO002/CoVispecT to P.B. and A.S. This research was partially supported by R01AI136514 (P.G.T).

## Author contributions

Conceptualization: E.R, M.V.P, A.A.M; Analysis: E.R, M.V.P, A.A.M.; Visualization: A.A.M, E.R; Resources: P.T, P.B., A.S, A.F.; Supervision and coordination: P.B. and P.T. Writing original draft: E.R, M.V.P., A.A.M. All authors provided discussion, participated in revising the manuscript, and agreed to the final version.

## Supplementary files

**Supplementary Table 1:** Differentially expressed genes for each functional Seurat cluster.

**Supplementary Table 2:** Unique alpha/beta TCR clonotypes enriched in COVID-19 cohort

**Supplementary Table 3:** Unique alpha/beta TCR clonotypes enriched in healthy control cohort

